# Cortical tracking of speakers’ spectral changes predicts selective listening

**DOI:** 10.1101/2024.05.23.595545

**Authors:** Francisco Cervantes Constantino, Ángel A. Caputi

## Abstract

A social scene is particularly informative when people are distinguishable. To understand somebody amid a ‘cocktail party’ chatter, we automatically index their voice. This ability is underpinned by parallel processing of vocal spectral contours from speech sounds, but it has not yet been established how this occurs in the brain’s cortex. We investigate single-trial neural tracking of slow frequency modulations in speech using electroencephalography. Participants briefly listened to unfamiliar single speakers, and in addition, they performed a cocktail party comprehension task. Quantified through stimulus reconstruction methods, robust tracking was found in neural responses to slow (delta-theta range) modulations of frequency contours in the fourth and fifth formant band, equivalent to the 3.5–5 KHz audible range. Instantaneous frequency spacing (Δ*F*), which also yields indexical information from the vocal tract, was similarly decodable. Moreover, EEG evidence of listeners’ spectral tracking abilities predicted their chances of succeeding at selective listening when faced with two-speaker speech mixtures. In summary, the results indicate that the communicating brain can rely on locking of cortical rhythms to major changes led by upper resonances of the vocal tract. Their corresponding articulatory mechanics hence continuously issue a fundamental credential for listeners to target in real time.

---

When attending a cocktail party, you might want to know *who* to hear from before *what*. The vocal chord system produces a speaker’s characteristic pitch, instantaneously subject to time-varying resonance effects due to the ever-changing articulatory manoeuvres of the supraglottal vocal tract. The resulting family of formant patterns jointly serves to cue and index speech from its vocal source. In many species, auditory processing of pitch and formants facilitates intraspecific recognition. These spectral cues often signal important anatomical and social features, including age, sex, body size, reproductive fitness, and social status or significance; for instance, observers may infer the likely size of an unseen conspecific given the average spacing between the caller’s time-varying formants (Fitch 1997; Riede and Fitch 1999; Collins 2000; Reby and McComb 2003; Bruckert et al. 2006; Evans et al. 2006; Puts et al. 2007; Puts et al. 2011). The versatility suggests that such spectral channels are routinely harnessed to broadcast and receive caller identity credentials through auditory space—a feat predating by far the emergence of language.

What spectral contour dynamics are represented through tracking activity by the auditory cortex or higher associated areas? Cortical tracking is the phenomenon describing the ability of network activity in the cerebral cortex to lock to the structure time-varying stimulus features as in speech, in which case it often refers to following its slow envelope. Such ability has in some cases been shown to correlate with listeners’ behavioral intelligibility levels (Vanthornhout et al. 2018). Analysis of cortical tracking while applying transcranial alternating current stimulation has further demonstrated the causal influence of phase-locked responses on intelligible speech perception (Wilsch et al. 2018; Zoefel et al. 2020).

Speech signals that are being attended to amid ambient noise elicit selective tracking responses, whereby the slow dynamics of cortical activity appears more consistent with the target speech envelope than the background (Ding & Simon 2012; Zion Golumbic et al. 2013; O’Sullivan et al. 2015). In scenes, selective tracking may manifest the existence of a cleaned-up representation of the target speech, where the accuracy of such representation possibly determines comprehension (Brodbeck and Simon 2020). There are now indications that cortical tracking, despite showing dominant responses to the speech signal’s envelope, also extends to some of the spectral cues of speech needed to establish speaker identity (Tang et al. 2017; Teoh et al. 2019; Van Canneyt et al. 2021; Brodbeck and Simon 2022). These studies have shown that the auditory cortex may track the dynamic contours and changes of voiced pitch, which offers opportunities to evaluate and generalize the association between cortical tracking and speech intelligibility. Much of speech intelligibility research has traditionally required systematic manipulation on spectral uncertainty, such as in noise-vocoded speech. In this regard, it is important to note that when the level of spectral detail is degraded to the point of unintelligibility, cortical tracking may be affected in spite of a preserved temporal envelope (Peelle et al. 2013). This underscores the need the contribution to tracking from the process resolving spectral contours in detail, as it offers a potential to connect with research demonstrate the tracking of higher-level linguistic units (Di Liberto et al. 2015).

Moreover, building on this area may offer an attractive way to probe the relationship of tracking frequency modulations generated by vocal tract resonances with behavior in the face of multiple voiced maskers. It specifically involves the question of what speaker-specific spectral features activate speech-tracking patterns in the brain that are relevant for speaker-selective behavior. Evidence of cortical encoding of frequency modulations (FM) in the voice hence represents a promising step to unravel how the brain targets “who” speaks under such conditions (Brodbeck and Simon 2022). A natural limitation in studies of spectral contour tracking is of course that cortical auditory population responses weakly resolve frequency-following responses even at pitch range (85-200 Hz). Yet, it is clear that representing spectral content at the higher frequency formant bands matters for cortical systems that establish speaker identity and phonemic analyses (López et al. 2013; Mathias and von Kriegstein 2014; Lee et al. 2019). We hypothesized that delta and theta band population activity in the cortex, part of which is known to track slow oscillations of the acoustic envelope of speech, may also provide a channel to represent additional slow changes of instantaneous FM in the formant range. To evaluate this possibility, we employed a stimulus reconstruction approach (Mesgarani 2014; Van Canneyt et al. 2021) designed to estimate the accuracy of recovering the variations of a given feature, such as instantaneous frequency, from a linear mapping of a listener’s electroencephalography (EEG) dataset. As with forward-model approaches (Teoh et al. 2019; Van Canneyt et al. 2021; Brodbeck and Simon 2022), decoding methods are evaluated in terms of their generalization performance. Due to scalp-wide, fine-grained multivariate information integration, backward modeling is a reliable means to test generalization performance. Performance from decoding models of envelope-tracking responses has been systematically used to predict behavioral responses in speech processing as objective measures of speech intelligibility (Vanthornhout et al., 2018). Importantly, when decoders show that specific stimulus information contributes to successful behavior, they effectively model the computational process that generates the observer’s expected response from coded stimulus features (Kriegeskorte and Douglas 2019).

In the present study, we addressed cortical tracking of spectral features in speech via reconstruction methods, at different bands across the formant range. Because of its potential basis for grouping features by speaker, we further approached the spectral feature which describes the instantaneous spacing pattern between formants arising as a result of their dynamic resonant structure. The average inter-formant spread has sometimes been used as proxy estimator for vocal tract length (VTL), a relevant parameter in automatic speaker recognition and vowel normalization among multiple talkers (Johnson 2020; Johnson and Sjerps 2021). Consequently, accurate decoding of a family of spectral signatures from the EEG of listeners, quantified through a single decodability score, may reasonably indicate their individual ability to follow dynamics stemming from changes in the apparent VTL of an unfamiliar talker as they speak. If reliable spectral contour reconstruction is quantified, such parameter may be furthermore expected to predict voice-specific selective listening ability of different listeners. We evaluated both hypotheses by, first, measuring the ability to faithfully represent spectral changes of different talkers in a single-speaker task through decodability scores. These were then compared with the behavioral accuracy of the same subjects probed on a comprehension report on aspects of target speakers’ identities, words and topics, as part of a selective listening task in a ‘cocktail-party’ scenario.

## Materials and Methods

### Participants

Seventy-three subjects (51 female; mean age 25.6 ± 4.9 SD) participated in the studies. Participants were native Rioplatense Spanish speakers with no hearing impairment or neurological disorder, and correct vision. Behavioral data from one participant was not stored in error. The Ethics in Research Committee at the Facultad de Psicología, Universidad de la República Uruguay, approved the study. All participants gave written consent in accordance with Declaration of Helsinki guidelines. Participants received a chocolate bar in appreciation for their time after they completed the task.

### Stimuli

Stimuli were obtained from original databases containing short phrase audios (mean duration 8.6 s, ±0.8 s) individually narrated by 607 speakers including 264 different male voices. The stimuli were sourced from Uruguayan news broadcasts, audiobooks, interviews, and recordings produced from Uruguayan volunteers. Recorded volunteer audios were read after phrases translated from a public database (Dolan and Brockett, 2005).

### Task

As part of the EEG study, participants were instructed to listen to each sound presentation with their eyes closed. In each trial, participants were first presented with a single-speaker speech stimulus (“single speaker listening”, SSL phase), followed by a cocktail party mixture stimulus in the second part (Supplementary Figure 1). Listeners were tasked with selectively attending to one of the two speech streams in the mixture, i.e., to select the leading or the interjecting talker. The indicated target stream was instructed randomly with equal probability. At the end of the mixture, participants were presented with a nine-option questionnaire. They were required to respond by exclusively selecting three options or more corresponding to the indicated target. Subsequent to response validation, a feedback screen was displayed with the trial and the accumulated scores (Supplementary Materials and Methods).

### Speech spectral feature analyses

Single-speaker speech stimuli from the SSL trial phase were submitted to an AM-FM analysis (Potamianos and Maragos 1996; Sadjadi and Hansen 2015; Mowlaee et al. 2016) in the original sampling rate. To optimize signal-to-noise ratio, the analysis was limited to male voice presentations as higher formants contain substantially less energy in female voices (Baumann and Belin 2010). Moreover, evidence of *f*_0_ tracking in the EEG to female voices is weak as decoder performance may be inversely proportional to frequency (Van Canneyt et al. 2021). A fixed 6-band gammatone filterbank was hence applied to each audio sample, assessing spectral variations corresponding to male *f*_0_ and formants’ *F*_1_-*F*_5_ frequency bands. Filterbank coefficients were based on six centre frequencies *fc_i_* set at 146, 671, 1870, 2870, 3946, and 4869 Hz that were defined from formant analysis of a local public voice database (López et al. 2013) conducted with the Praat software. Coefficients were estimated with the Large Time/Frequency Analysis toolbox (Søndergaard et al. 2012; Průša et al. 2014). Channel bandwidths equalled thrice the critical bandwidth of the auditory filter at the corresponding *fc_i_*, and were defined in equivalent rectangular bandwidths (ERBs) (Glasberg and Moore 1990). The real part of the filter coefficients was applied to each audio signal. Each filter output was analyzed to obtain its instantaneous frequency *f_i_*(*t*), which was estimated by differentiating the instantaneous unwrapped phase *Φ_i_*(*t*) of the corresponding analytic signal, computed by the Hilbert method. Expressed in polar coordinates, the analytic signal conveniently separates from the effects of instantaneous amplitude content *a_i_*(*t*), which is extracted in envelope tracking for instance. A multiband demodulation analysis was performed by mean-amplitude weighted short-time estimation of the instantaneous frequency per sub-band (Grimaldi and Cummins 2008). At time instant *t*_0_, the short-time estimate for the spectral modulations in the frequency *F_i_*was obtained by integration of the weighing instantaneous amplitude function over a τ=25 ms window, according to the formula:

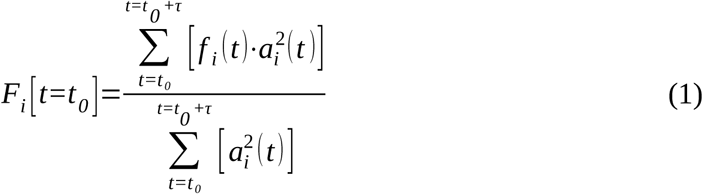

In effect, spectral modulations are separated from amplitude modulations within each frequency band by rendering the band’s estimated instantaneous frequency first, and smoothing it over short windows, weighing each frequency by the relative power of the filtered signal at that specific instant. The resulting set of discrete short-time weighted instantaneous frequency estimates *F_i_*(*t*) is sometimes referred to as the pyknogram of the speech signal (Grimaldi and Cummins 2008). It may be used as a basis for speech formant estimation, alternatively to linear predictive coding methods for instance (Potamianos and Maragos 1996). Here, the pyknogram was used to represent the dynamic waveform timeseries for instantaneous frequencies within the respective ranges of *f*_0_ and formants’ *F*_1_-*F*_5_.

The dynamic waveform for the instantaneous frequency spacing Δ*F* measure was constructed from *k*=[1,…,*K*] successive independent linear regression models, with *K* the total of time samples in the stimulus. At the time instant *t_k_* represented by the *k*-th sample, the model of the spectral observation *F_i_* is:

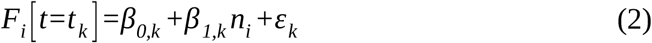

with β_1_ representing the linear slope of the regression model of *F_i_* values against their associated formant band numbers *n_i_*={1,2,3,4,5} (the model predictors), and β_0_ the intercept; ε represents the linear model error. Model coefficient estimation was performed via an iteratively reweighted least squares algorithm with the bisquare weighting function (MATLAB `robustfit`). The regression slope served as the measure of spectral spacing (Johnson 2020):

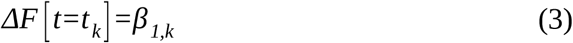

The seven resulting spectral contour timeseries waveforms (*f*_0_, *F*_1_*–F*_5_, Δ*F*) were downsampled to 1024 Hz.

### EEG preprocessing

EEG data analyses were implemented in MATLAB 2018b. EEG signals were bandpass-filtered between 1 and 28 Hz with a 10-order elliptic IIR filter of 60 dB attenuation and 0.5 dB passband ripple, with group delay correction. Single channel data were rejected by a channel variance-based criterion (Junghöfer et al. 2000) blind procedure, implemented with confidence coefficient *λ_P_*=4 . The procedure was performed separately for the external reference channels, with a more restrictive confidence coefficient *λ_P_*=2 . Head (and reference) sensor datasets were common median-referenced, after which the automated rejection procedure was performed a second time on the full set. EEG data were decomposed into independent components using FastICA (Hyvarinen 1999) and the two independent components with maximal broadband power were automatically selected and projected out of the data. A time-shifted principal component analysis (de Cheveigné and Simon 2007) was then applied to discard environmental signals recorded on the external reference sensors, using ±4 ms time shifts. A sensor noise suppression algorithm was used to attenuate artifacts specific to any single channel, using 63 neighbors (de Cheveigné and Simon 2008).

EEG data time samples with a value greater than 7 standard deviations were set to zero after which single trial neural data were aligned with stimuli data, and trials were concatenated in time. The set of EEG and stimulus signals was then band-pass filtered between 1 and 8 Hz with a second-order Butterworth filter in the forward and reverse direction, downsampled to 128 Hz, and converted to standardized units.

### Stimulus-response alignment and decoder estimation

Using EEG responses from single speaker listening phases, we assessed performance in reconstructing the slow modulations of the instantaneous frequency waveform of speech at (*i*) the *f*_0_ band, (*ii-vi*) the first to fifth formant bands (*F*1 to *F*5), (*vii*) the spectral spacing parameter Δ*F*; and (*viii*) the instantaneous waveform of speech broadband envelope (see *Supplementary material*); resulting in eight separate decoders based on the empirical EEG activity, per listener. Eight additional decoders were separately estimated, based on trial-reshuffled EEG activity, so as to compare each individual empirical performance measure to an estimation noise reference.

The operation to obtain the linear reconstruction *S̃_i_*(*t*) of a univariate stimulus timeseries *S_i_*(*t*) from a set of *N* EEG channel responses *R_n_*(*t*) can be formulated as:

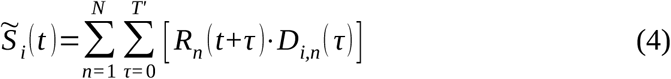

where *D_i,n_*(*t*) is the *i*-th linear decoder that operates on the response timeseries *R_n_*(*t*) from channel *n* in order to reconstruct the *i*-th acoustic feature. Each individual decoder *D_i_* is a *N* ╳ *T’* matrix with *N* = 64 channels and *T’* = 400 ms denoting the size of the temporal integration window. Hence, at time point *t*=*t_k_*, the stimulus timeseries is reconstructed by linear combination of all channel responses spanning from *t_k_* until *t_k_*+0.4 s. Decoders were obtained using the boosting method with 10-fold cross-validation (David et al. 2007; Puvvada and Simon 2017). The resulting stimulus reconstruction approximates the true stimulus by a residual response waveform *ɛ_i_*(*t*) not explained by the linear decoder model:

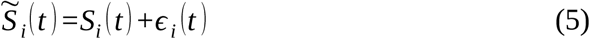

Empirical decodability of the *i*-th feature, Π*_i,e_*, is measured by the Spearman rank correlation between *S_i_*(*t*) and *S̃_i_*(*t*). For each feature we additionally obtained its reference decodability Π*_i,r_*, defined as the Spearman rank correlation between *S_i_*(*t*) and the linear reconstruction that results from training a decoder *D_i_*\* based on trial-reshuffled EEG data *R_n_*\*(*t*). Decoder reliability (also referred to as neural decoding performance here), was expressed as _(_*ρ_i,e_*− *ρ_i,r_* _)_/ *ρ_i,r_* *100%, i.e., the percent relative change of empirical decodability with respect to the performance of the corresponding reference decoder.

### Decoder performance as predictor of listening performance at the cocktail party

Upon identification that the reliably decodable spectral features were the modulation from bands *F*_4_, *F*_5_, and the Δ*F* measure (see Results), their common empirical decodabilities were indexed by the Euclidean norm, per participant:

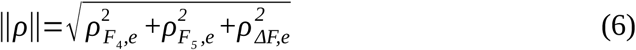

A participant’s high score in this more general spectral feature reconstruction parameter represents systematically higher decodability of all three significant spectral features. ||*Π*|| was used to evaluate the hypothesis that cortical tracking of vocal tract features from speech predicts listener performance upon facing cocktail-party scenarios. The proportion of correct trials in the cocktail party task was taken as a measure of participant behavioral performance. The association between subjects’ ||*Π*|| and cocktail-party performance was evaluated with Spearman’s rank correlation *r_s_* with interval testing (Pernet et al. 2013), and confirmatory tests were subsequently performed via Shepherd’s correlation (Schwarzkopf et al. 2012) and robust regression with the Huber weighting function.

## Results

The ability to decode seven acoustic features that relate frequency modulations in voiced speech, in addition to the speech envelope, was investigated from EEG recordings using stimulus reconstruction methods. The spectral features are delta-theta band modulations of instantaneous frequency modulations in (*i*) the fundamental frequency band, (*ii*-*vi*) the *F*_1_, *F_2_*, *F_3_*, *F_4_*, *F_5_* formant bands, and (*vii*) their interband spacing (Figure 1). Decoder analyses were based on single-trial, single-speaker male voice listening epochs, as in the EEG these offer relatively higher precision frequency tracking data than female speaker listening [9]. Figure 2A depicts an example of stimulus reconstruction applied to delta-theta band modulations of the speech envelope from one sentence stimulus. Here, the original envelope waveform is reconstructed from a decoder based on empirical EEG data to a degree that, in relative terms, exceeds what a second, independent decoder may achieve using reshuffled EEG data as reference. In a similar way, the same EEG data additionally contain tracking responses to the dynamics of instantaneous interband frequency spacing Δ*F*, which denotes the time-varying dispersion of spectral resonances in a speaker’s voice.

**Figure 1.**
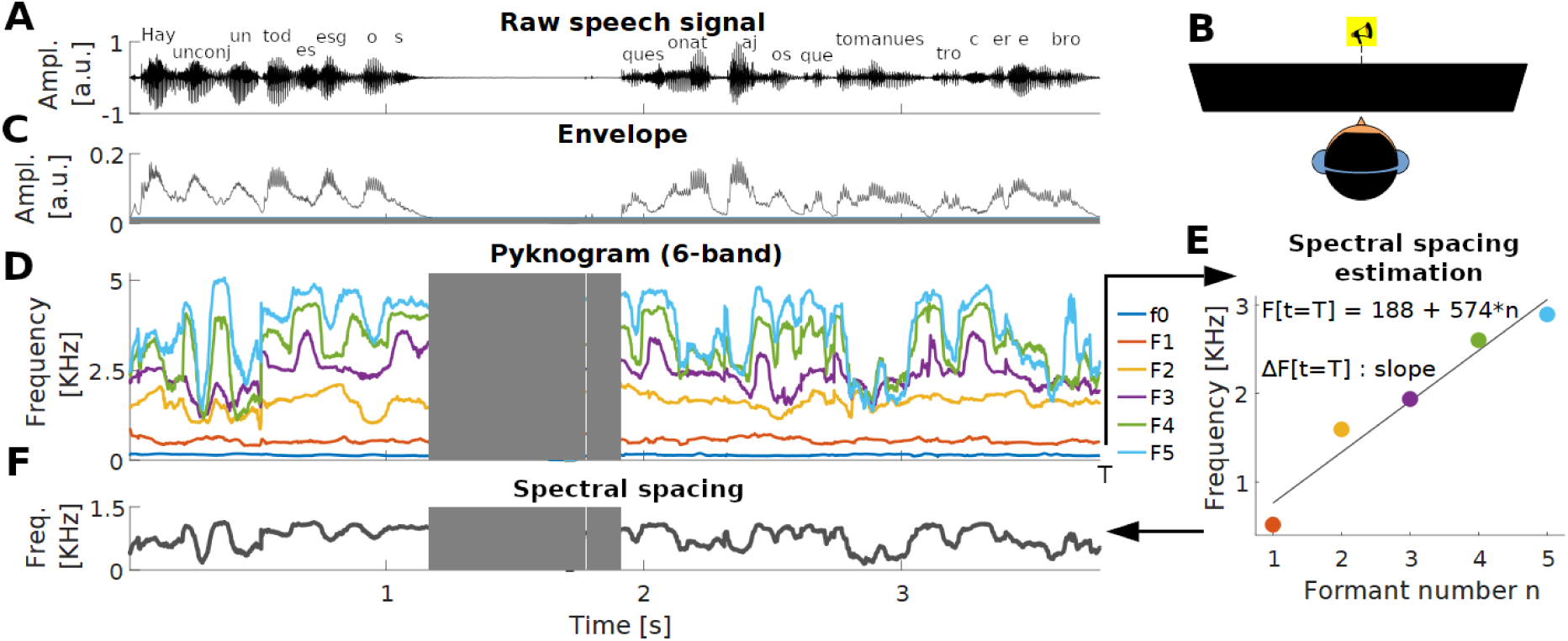
Setup for extraction of spectral features from voiced speech. **(A)** Trial of a single-speaker speech sentence. **(B)** Participants listen diotically to centrally presented speech trials while undergoing EEG. **(C)** Broadband envelope of the speech sentence. **(D)** Instantaneous frequency estimates from the sentence. Each estimate corresponds to a spectral band centred on the fundamental frequency *f*_0_ or *F*_1_-*F*_5_ formant bands. Silent regions (grey areas) are not analyzed. **(E)** Method of Δ*F* estimation, based on linear regression of instantaneous frequency values indexed by the number of the associated formant band. For this example, Δ*F* at time sample *T* is 574 Hz.

**Figure 2.**
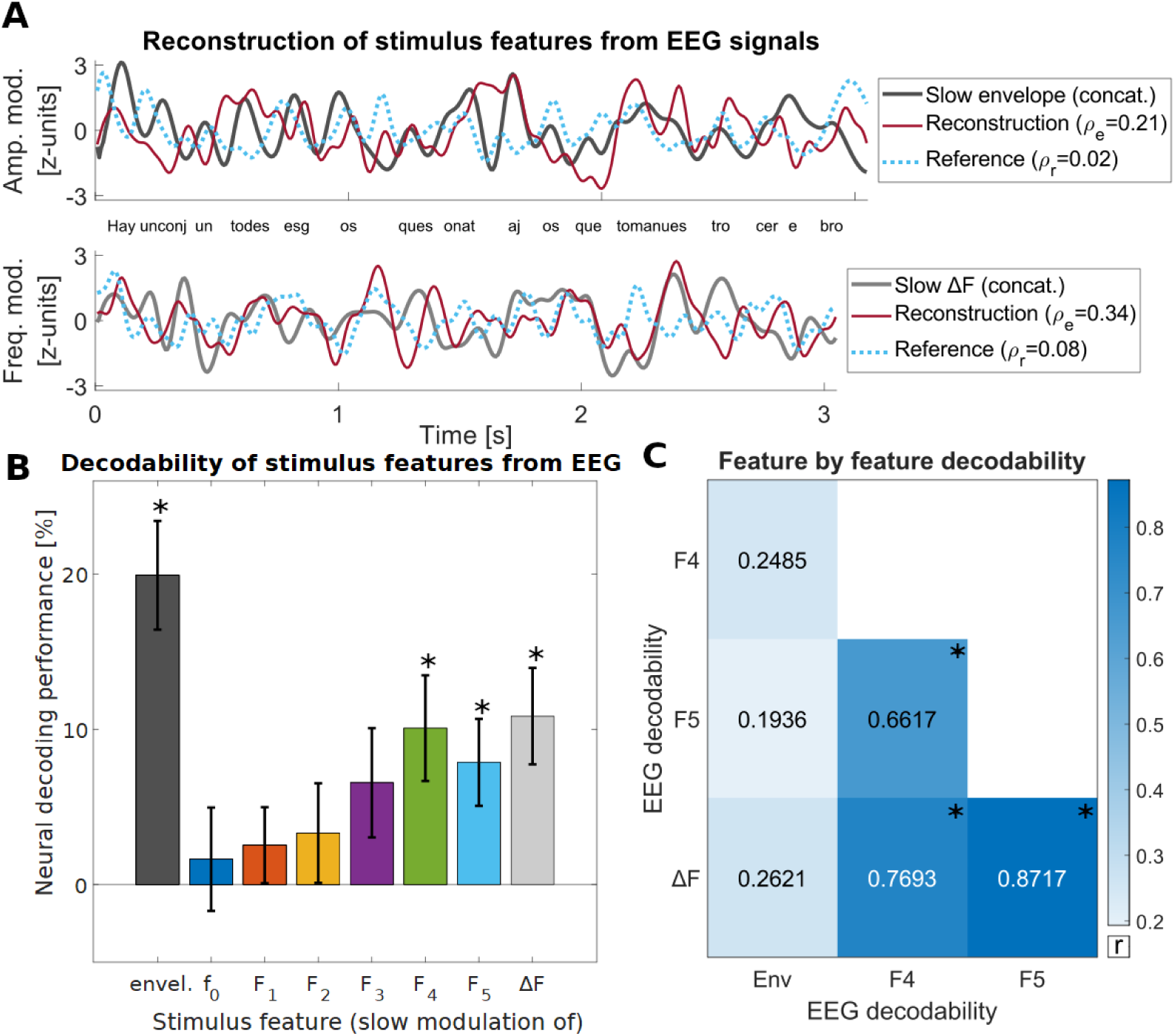
Spectral feature waveforms from voiced speech are reliably decoded from single-trial EEG. **(A)** Sample reconstruction of speech feature waveforms after the sentence example from Figure 1. Two estimated decoders per listener (*N* = 19 for this stimulus) were obtained, corresponding to empirical versus reshuffled EEG datasets. *Top*: Slow modulations of the broadband envelope, plus subject-averaged envelope reconstructions by decoder type. Reconstruction performance is here represented by the rank correlation between the stimulus and mean reconstruction; the empirical performance exceeds that of reference decoders estimated after reshuffled EEG data. *Bottom*: Application of the method to a spectral vocal feature, instantaneous frequency interband spacing (Δ*F*). **(B)** The slow amplitude modulations of the envelope, frequency modulations of spectral features from the higher formant range, and β*F,* are reliably decoded from single-trial EEG across listeners. **(C)** Pairwise correlations between acoustic feature decodabilities, across listeners. The three robustly decodable spectral features significantly correlate with each other, indicating that neural tracking across these dimensions is mutually predictable. These decodabilities however do not predict decodability of the amplitude dynamics of the broadband envelope.

### Robust decodability of vocal spectral dynamics from single-trial EEG recordings

Neural decoding performance, quantified by the empirical-to-reference reconstruction performance ratio, referred to as empirical Π change, was estimated per acoustic feature (Figure 2B). The slow (i.e., delta-theta) modulations of the broadband envelope were, as expected, robustly decoded from the single-trial EEG (mean empirical Π change 19.9%, SD = 29.9%) *t*(72) = 5.7, adjusted (Bonferroni-Holm) *p* = 10^-5^. In addition, the analyses showed that slow modulations of the instantaneous frequency waveforms from higher formant ranges were also reliably decodable: *F*_4_ (mean 10.1%, SD=29.1%) *t*(72) = 3.0, adj. *p* = 0.013; *F*_5_ (mean 7.9%, SD=23.8%) *t*(71) = 2.8, adj. *p* = 0.016. Furthermore, robust decodability was also found in EEG responses to the dynamics of instantaneous interband frequency spacing Δ*F* (mean 10.8%, SD = 26.5%) *t*(72) = 3.5, adj. *p* = 0.003. Supplementary Figure 2 presents the raw (empirical) reconstruction performance in terms of Spearman rank correlations.

At the lower frequency ranges, decodability of slow modulations of the instantaneous frequency waveform from the fundamental *f*_0_ band was not found to be robust (mean 1.6%, SD = 28.4%) *t*(72) = 0.5, adj. *p* > 0.31. Null results were similarly observed for decodabilities of spectral features from: *F*_1_ (mean 2.5%, SD = 20.9%) *t*(72) = 1.0, adj. *p* > 0.15; *F*_2_ (mean 3.3%, SD = 27.4%) *t*(72) = 1.0, adj. *p* > 0.15; or *F*_3_ bands (mean 6.6%, SD=30.1%) *t*(72) = 1.9, adj. *p* = 0.13. Hence, slow modulations in the spectral variations across the lower frequency bands of voiced speech were not reliably decoded from single-trial EEG data. The results indicate that the dynamics of spectral contours associated with the higher formant ranges, and spacing information used to index vocal identity, may be however tracked by cortical populations during natural speech listening.

We next addressed whether listeners that achieve better envelope decodability similarly achieve higher decodabilities across the relevant spectral domains *F*_4_, *F*_5_, and Δ*F*. Using a Pearson correlation matrix on subject decodability data by feature (Figure 2C), significant linear relationships among feature pairs were found, although these were limited within all spectral feature pairs (all adj. *p*-values < 1.6×10^-4^). Subject decodability for any of these features did not predict subject decodability for the envelope (all adj. *p*-values > 0.6). These results indicate that listeners may encode the spectral features of speech by relying on temporally matching modulations across spectral variations and their spacing patterns in the formant range. In addition, the results suggest a distinct acoustic processing pathway from that which encodes sound level variations signaled by the speech broadband envelope.

### Spectral tracking abilities predict performance gains at the cocktail party

Facing new, unfamiliar speakers, selective listening depends critically on the accurate delimiting of spectral sets that target a speaker’s content. Our second aim was to test whether a listener’s spectral tracking ability may help her select and resolve speech from a cocktail party of unfamiliar speakers. Spectral tracking ability was quantified by ||Π||, the L2-norm of the vector of empirical decoder performances (Π*_e,F_*_4_, Π*_e,F_*_5_, Π*_e,_*_Δ*F*_) which increases as a subject consistently encodes all three relevant spectral features. We hypothesized that higher ||Π|| may boost a listener’s chances to reliably track a masked speaker of choice, as participants also performed a ‘cocktail-party’ comprehension task. Their level of intelligibility during this stage of the experiment was evaluated through a challenging comprehension questionnaire (see *Supplementary Materials and Methods*). The proportion of correct cocktail party trials that were achieved by the listener was used as index of performance. A robust correlation analysis with interval testing (Pernet et al. 2013) showed that the summary neural measure ||Π|| and behavioral cocktail party task performance were found to be positively correlated, *r_s_* = 0.39, CI = 0.18 – 0.57, *p* = 8.05×10^-4^ (Figure 3A). The significant correlation was robust to potential outliers (Shepherd’s Pi = 0.41; *p* = 0.001), and confirmed by a robust regression analysis *t*(70) = 2.994; *p* = 0.004. Furthermore, ||Π|| similarly predicted behavioral data when expressed in terms of the amount of points earned per trial, *r_s_* = 0.36, CI = 0.13 – 0.55, *p* = 0.002 (Figure 3B), confirmed by robust regression *t*(70) = 3.032; *p* = 0.003.

**Figure 3.**
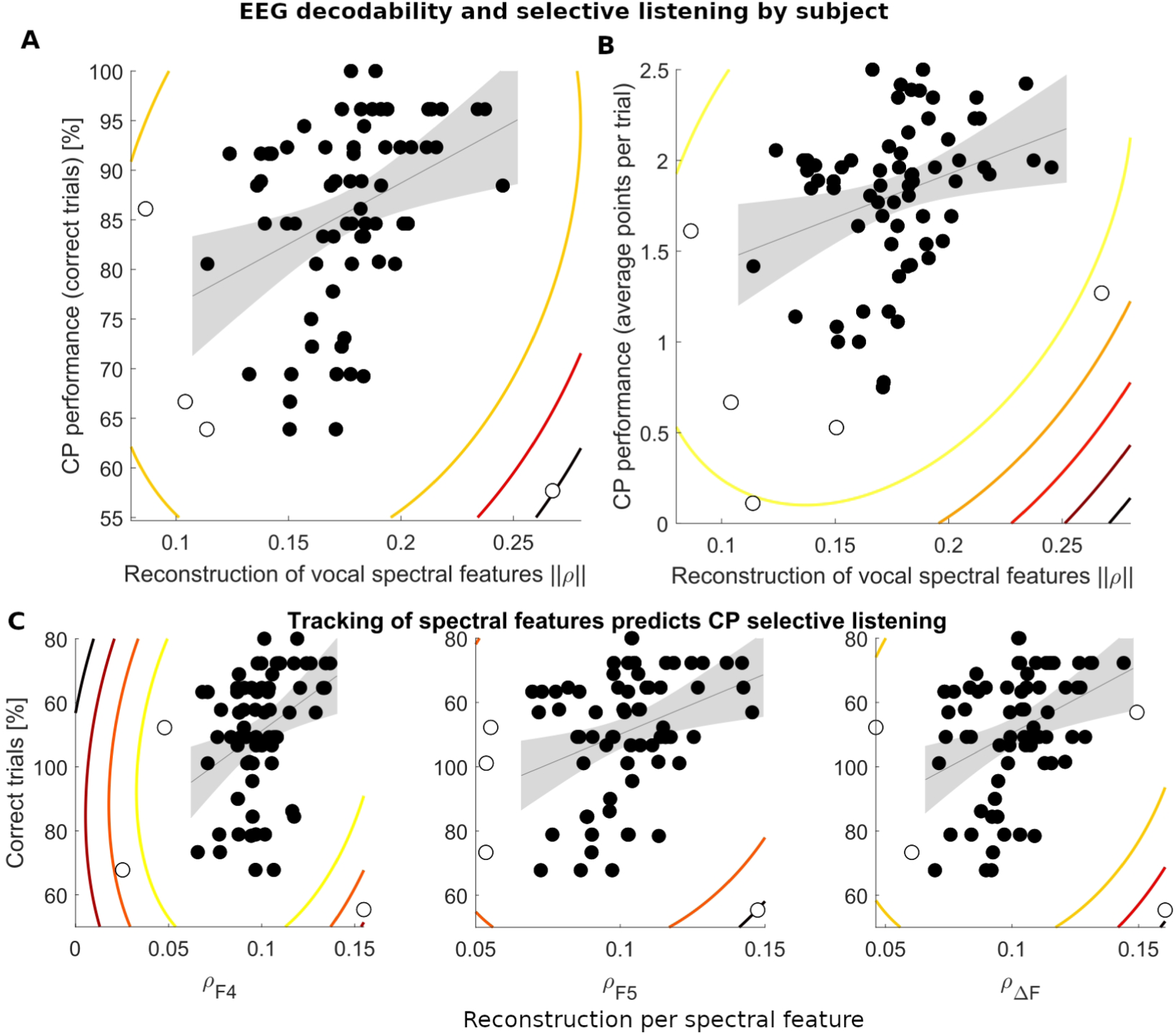
Spectral features of voiced speech assist performance in multispeaker conditions. **(A)** A summary measure of neural decodability across the three relevant spectral features significantly predicts the behavioral performance of listeners at the ‘cocktail party’. **(B)** Joint decodability of spectral contours also predicts subject performance measured by mean points per trial at the cocktail party task, with a significant robust correlation relationship observed, Shepherd’s Pi = 0.32, *p* = 0.015. **(C)** Decodabilities of higher formant range instantaneous frequencies (*F*_4_, *F*_5_) and instantaneous frequency spacing (Δ*F*) dynamics individually predict performance at the cocktail party task. Robust correlation relationships are observed for *F*_4_ (left), Shepherd’s Pi = 0.39, *p* = 0.002; *F*_5_ (middle), Shepherd’s Pi = 0.35, *p* = 0.007; and Δ*F* (right), Shepherd’s Pi = 0.37, *p* = 0.004. Contour lines indicate distance *D*_s_ from the bivariate distribution mean, expressed in z-units (squared) and shown from *D*_s_=12 in steps of 6 (Schwarzkopf et al., 2012). Points with *D*_s_ ≥ 6 are defined as outliers.

The cocktail party questionnaire includes items probing three different aspects of target speech, namely, the speaker’s vocal identity, the keywords in speech, and the classification of its subject matter or topic (see Supplementary Methods). The ability of ||Π|| to predict responses in each of these domains separately was further analyzed. First, ||Π|| was found to predict the likelihood of listeners’ correct choices about target speakers’ identities, *r_s_* = 0.31, CI = 0.09 – 0.51, *p* = 0.007 (Supplementary Figure 3A), confirmed by robust regression *t*(70) = 3.225; *p* = 0.002. Second, ||Π|| was similarly found to predict the likelihood of listeners’ correct choices about target speech keywords, *r_s_* = 0.32, CI = 0.09 – 0.54, *p* = 0.006 (Supplementary Figure 3B), confirmed by robust regression *t*(70) = 2.382; *p* = 0.020. Third, ||Π|| did not significantly predict the likelihood of listeners’ correct choices about target speech topic, however, *r_s_*= 0.21, CI = -0.03 – 0.41, *p* = 0.078 (Supplementary Figure 3C). Moreover, cocktail-party data involved voices of either sex, and we verified by robust regression that ||*Π*|| measures of spectral tracking from male listening EEG data also predicted listener’s cocktail party behavioral perfomance during the targeting of a male speaker, *t*(70) = 2.913; *p* = 0.004. The positive correlation similarly held for the subset of cocktail party trials exclusively involving male+male voice combinations, *t*(70) = 3.192; *p* = 0.002.

Finally, we examined whether the neural tracking of spectral features (Π*_e,F_*_4_, Π*_e,F_*_5_, Π*_e,_*_Δ*F*_) may individually predict cocktail party performance (Figure 3C). In each case, the individual neural tracking measure and behavioral cocktail party task performance were found to be positively correlated, i.e., *F*_4_: *r_s_* = 0.35, CI = 0.11 – 0.55, *p* = 0.002; *F*_5_: *r_s_* = 0.32, CI = 0.09 – 0.53, *p* = 0.006; Δ*F*: *r_s_* = 0.32, CI = 0.09 – 0.51, *p* = 0.006. Each of the significant correlations was confirmed by robust regression analysis, *F*_4_: *t*(70)=3.259, *p*=0.002; *F*_5_: *t*(70)=2.587, *p*=0.012; Δ*F*: *t*(70)=2.563, *p*=0.013. The findings are consistent with the notion that successful selection in multispeaker conditions is promoted when cortical responses follow through major changes of spectral attributes that serve to distinguish between vocal tracts.

## Discussion

Cortical systems that process natural speech avail themselves of higher-range frequency modulations, as demonstrated here with noninvasive single-trial EEG methods. First, stimulus reconstruction techniques retrieved, as they do in the case of the speech envelope, delta-theta band modulations contained in various spectral indices of speaker identity. Subsequently, listeners’ ability to reconstruct spectral features predicted success at following a talker’s speech in a ‘cocktail-party’. Slow modulations to instantaneous spectral variations from higher formant bands *F*_4_ and *F*_5_ were reliably decoded from EEG, yet this ability was not explained by reconstruction of amplitude modulations contained in the envelope, suggesting alternate processing mechanisms. Decoding findings were also presented here for a key feature in voice recognition studies, the spacing of instantaneous frequencies between formant bands (Δ*F*), which may be relevant because inter-formant dispersion is often used to infer anatomical dimensions of the source’s vocal tract (Lavan, Knight, et al. 2019), (Lammert and Narayanan 2015). All three spectral features considered, their joint decodabilities predicted listeners’ ability to selectively attend in a two-speaker scenario.

### Cortical tracking of spectral contours in the higher formant band

The peripheral auditory system isolates frequency-localized components of speech and subsequent central processing for auditory scene analysis of these components occurs mainly in the time domain (Friedman 1985). Extracted pitch and spectral cues, including formants, serve to build representations of a speaker’s voice as listeners face the problem of determining speaker invariance (Lavan, Knight, et al. 2019). This is a crucial aspect of listeners’ organization of multispeaker scenes, which needs estimating the variability that corresponds to one voice with sufficient precision in order to set it apart from maskers’ (Darwin et al. 2003) (Lavan, Burton, et al. 2019). Studies suggest that the first 400 ms of auditory processing perform the necessary spectral vocal cue processing analyses (Levy et al. 2001; Charest et al. 2009; Capilla et al. 2013). Continuous sequential analyses of the unfamiliar voice signal are traditionally assumed to trigger the computational process of perceptual voice-identity processing where identity-features are extracted and merged, creating a coherent voice percept (Maguinness et al. 2018). Here, we present sequential filterbank analyses of a wide collection of voice stimuli, based on the phase of the time-varying analytic signal at each band. The results demonstrate cortical tracking coding of frequency (rather than amplitude) modulations within the formant range. It is important to consider the fact that the stimulus reconstructions are based on time-varying frequency contours of content within formant bands, without this necessarily be equal to the formants themselves. The present findings therefore build on previous studies of pitch contours tracked at the auditory cortex (Van Canneyt et al. 2021; Teoh et al. 2019; Brodbeck and Simon 2022). With regards to inconsistent findings on *f*_0_ decoding between these prior reports and the current study, we note three important classes of differences. The first relates to differences in the stimulus representation, where broadband rather than low-frequency temporal modulations of pitch waveforms were previously addressed, assessing spectral variations at correspondingly finer temporal resolutions. In the case of Van Canneyt and colleagues, evidence of *f*_0_ tracking was most robust at the brainstem stage (Etard et al. 2019) but cortical responses up to 40 ms were also inspected. They found cortical tracking to be optimal for male speaker stimuli voiced with low, steady *f*_0_ values. In our case, we found spectral cue decoding across the range of male speakers comprising middle and late stage activity from a 400 ms period, but this did not include the *f*_0_ band. The inconsistency would tentatively suggest that phase-locked responses to the coarser, slow modulations of *f*_0_ that we sample may vanish from population activity following a temporal code, over cortical stages across multiple speakers.

Second, both Teoh et al., 2019 and Brodbeck and Simon, 2022 focussed on cortical responses up to 300 ms (or more), finding cortical tracking of *f*_0_ contours in the low frequency bands of M/EEG responses. Unlike Van Canneyt et al., 2021, the stimuli representations they investigated convey critical information about all silent periods in presented speech. For this reason, their findings of *f*_0_ tracking responses may be potentially accompanied by on/off response coding that is also signaled via envelope tracking. We consider that such types of contribution to forward models of any continuous stimulus feature may result in small but non-negligible performance advantages over distributions of trial-reshuffled data. For this reason we recommend that, given the dominance of envelope tracking in auditory cortical responses, silent periods be removed prior to modeling whether in *f*_0_ tracking or other standalone continuous features (an alternative is to introduce a voicing binary variable, cf. Tang et al. 2017). When envelope tracking is compared against tracking of other features as is the case here, it is further adequate to excise unvoiced periods from the envelope signal itself as well. Despite remaining segments missing all elements of natural prosody (of which silences may play an important role), removal provides for comparisons of resolving spectral contours unbiased by systematic temporal edge responses.

Third, prior studies (Teoh et al. 2019; Van Canneyt et al. 2021; Brodbeck and Simon 2022) addressed coding models of a speaker’s normalized relative pitch. By contrast, our backward decoding models incorporate joint data across multiple speakers on a common spectral frame. The present approach hence rather models the temporal modulations of absolute changes in *f*_0_ waveforms across speakers because they are proportional to their differences. Using electrocorticography, Tang et al. 2017 directly addressed the distinction between absolute and relative pitch encoding in cortical networks. Their findings indicated the two types of encoding may be subserved by separate networks in the auditory cortex, with a possible dominance of relative pitch encoding. Our negative finding might reflect that network activity from absolute pitch encoding does not generally predict neural discriminability of temporal responses to different intonation contours in the auditory cortex (Tang et al. 2017). Although the previous M/EEG studies employed one or few different speakers, their *f*_0_ tracking findings appear consistent with relative pitch encoding, which does predict neural discriminability of intonation contours. Nevertheless, from the perspective of a ’cocktail-party’ setting, tracking models of absolute spectral features, including the formant range, remain crucial because they are the basis for speaker voice segregation. Our lack of significant findings for spectral cues in the *f*_0_, *F*_1_–*F*_3_ bands may suggest a lower bandwidth limit for slow absolute FM changes encoded in delta/theta activity. It is worth noting that, while phonemically important fast rates of change occur in movements of the lower formants that encode short vowels (Baumann and Belin 2010), higher formant band changes appear less systematically related to vowel quality. Moreover, vowel encoding is considered to be subject to subsequent speaker-invariant representations via rescaling processes (Johnson and Sjerps 2021). As such, determination of spectral cues from higher formant bands, in absolute terms, could provide a reliable channel for sustained discrimination of unfamiliar voices over the first few seconds of multi-speaker listening.

### Spectral tracking as predictor of listeners’ speech segregation performance

Spectral contour reconstruction analyses further demonstrated their ability to relate the likelihood of listeners’ correct choices on relevant aspects of comprehension behavior in a ’cocktail-party’ task. One fundamental aspect of effective speaker selectivity is the successful discrimination of target speaker identities from the background, such as on the basis of their perceived sex or age. Neural decodability measures predicted such ability, which is consistent with the encoding of absolute spectral features from the dynamic voice signal as a basis for correct speaker identification. Importantly, reconstruction accuracy analyses demonstrated the ability to predict the likelihood of listeners’ identification of correct word utterances from the target speaker. This finding is consistent with the involvement of spectral contour tracking in the processes that lead to extraction of lexical content during selective speech listening. At the level of topic information, we did not find evidence of similar correlations of listener’s correct choices with reconstruction accuracy however. This null result may reflect the limitations of relatively lower-level spectral cue coding in the inference of higher-level thematic content.

Neural measures obtained under optimal listening conditions remain a key element of models of speech intelligibility for sub-optimal listening conditions with irrelevant sources, such as different noise levels (e.g. neurometric lapse rate, Vanthornhout et al. 2018). Here, neural measures were obtained from the single-speaker task, and used to predict behavioral performance of the two-speaker cocktail-party task. Hence, we tested whether the quality of neural representations from noise-free scenarios is useful to predict the degree of listeners’ intelligibility in multiple-speaker listening conditions – without recourse to its likely contingency on noise adaptation mechanisms (Khalighinejad et al. 2019). This means that, in our behavioral testing, we did not specifically address whether neural representations from ideal conditions remain noise-invariant. Nevertheless, the observation that such representations impact behavioral performance may suggest that, across listeners, clean representations may be commensurate with those potentially achieved under noise adaptation, although this remains an open question.

For the reason above, our current approach may be insufficient when one aims to predict within-subject intelligibility across interference levels, for instance. Yet, it may be advantageous regarding the auditory scene analysis of elements continually subject to causal inference during multi-speaker masking. This refers to the process by which the observer adjudicates common or separate causes (e.g. speakers) to the auditory signal. Its computation may involve the different resolved spectral contours and their combinations, where clean speech occurrences from occurring masker-free periods become particularly useful. Prior research has shown that processing of such overt periods is expedited when part of the attended target, as opposed to its masked elements, whose processing is delayed (Brodbeck et al. 2020). In practical terms, responses based on single-speaker epochs convey listeners’ tracking ability of a target’s key spectral features for periods that are exempt from causal inference. Such opportunities represent a means for listeners to constrain and establish an unfamiliar vocal identity from the varying speech signal on summary statistical patterns (Lavan, Knight, et al. 2019). In computational voice coding models, static pitch and formant dispersion parameters have traditionally represented the coordinates of hypothesized vocal space (Baumann and Belin 2010; Latinus et al. 2013; Lavan, Knight, et al. 2019). These follow from distinguishable origins in source and filter models of production, namely, the individual contributions made by the larynx to pitch and by the supra-laryngeal vocal tract to formant structure. Spectral cues that are based on formant spacing may show higher reliability for speaker identification than pitch (Gaudrain et al. 2009). Inter-formant spacing is extracted by the auditory system (Ives et al. 2005; Smith et al. 2005; Pisanski et al. 2014) as early as at the medial geniculate body (von Kriegstein et al. 2006), and is cortically represented in posterior superior temporal lobe networks that include temporal voice areas (von Kriegstein et al. 2007; von Kriegstein et al. 2010; Mathias and von Kriegstein 2014). These regions may extend across the posterior aspect of the right superior temporal sulcus to the superior temporal pole (Belin et al. 2000; Belin and Zatorre 2003; Belin et al. 2004; Kriegstein and Giraud 2004; Latinus et al. 2011; Latinus et al. 2013; Pernet et al. 2015; Belin 2017; Belin et al. 2018; Mathias and von Kriegstein 2019). Responses in this region may be considered to be involved in low-level voice processing when they remain specific to voice-like stimuli regardless of intelligibility (Rosen et al. 2011). For this circuitry, multispeaker conditions is naturally impose short time spans to attempt reliable voice discrimination.

Spectral cues within these reduced windows then provide variations of an unfamiliar voice that may be harnessed and reliably summarized as in clean conditions. Albeit at a slower rate, the process could sequentially buildup to form within-person vocal prototype representation, e.g. (Lavan, Knight, et al. 2019; Lavan, Burton, et al. 2019). Future studies may determine whether tracking abilities, at masked periods, possibly sustain the likelihood that an inferred spectral grouping can be maintained in consideration that alternative formant complexes are part of causal inference.

Finally, many studies have consistently shown the neural tracking of more abstract and higher-level properties of speech than shown here, ranging from detailed narrowband amplitude dynamics to phonemic changes and lexical probabilistic structure (Di Liberto et al. 2015). The present work extends these prior findings by introducing a basis for effective targeting of a *caller* of choice, with potential selective communication across wider animal models. Lexical processing of speech is constrained by the precise encoding of vocal tract modulations (Perrachione and Wong 2007; Chandrasekaran et al. 2011; Perrachione et al. 2011; McGettigan and Scott 2012; Zäske et al. 2017). Relative temporal displacements between spectral components affect intelligibility performance (Stilp et al. 2010), for example from surface reverberation effects that lead to weaker segregation in multispeaker conditions (Abouchacra et al. 2011; Sayles et al. 2016). Bandwidth tuning loss at higher frequencies under presbycusis also severely limits the ability to target speech in multispeaker conditions. The process by which these levels of uncertainty affect the monitoring or tracking of spectral groupings and, more generally, the causal arbitration to integrate or segregate them, remain unclear. Future studies may also address what impact do limitations to tracking of spectral features of speech based on multiple formants at any given time, imply for normal hearing and clinical populations.

## Conclusion

The human voice is acoustically fingerprinted, thanks in part to the shape and function of the larynx and oral cavities as they change from person to person, and from one instant to the next. This study shows that a substantial portion of these time-varying signatures are consistently traced by cortical networks. People who are more capable of tracking these variations neurally are more likely to grasp speakers of their choice despite masking by others. Thus, the changes can be used for the brain to tell what pieces of an unfamiliar voice fit together and which ones to put apart. Solving this inference in real time represents a cornerstone to understanding communication in everyday’s social world.

## Acknowledgments and funding sources

Conflict of interest: None to declare. Original EEG and behavioral data were from Thaiz Sánchez-Costa’s Master’s thesis project, and we are thankful for her excellent data collection. We also thank Rodrigo Caramés Harcevnicow for his assistance in data analysis. The research presented in this manuscript received funding from the Agencia Nacional de Investigación e Innovación, Uruguay, under the research grant FCE_1_2019_1_155889 to FCC.

## Supplementary Materials and Methods

### General overview

The data upon which this article is based on is part of a larger dataset that was obtained in the course of a series of studies on the cocktail party effect (Sánchez-Costa et al., results to be published elsewhere). These original EEG studies aim to investigate how prior speech listening experience modulates speech segregation, based on forward modeling during multi-speaker listening. In the present study, we addressed the stimulus reconstruction (i.e., backward modeling) of acoustic features in single-speaker speech. Neural estimates obtained from these data were compared with behavioral performance data at multi-speaker listening (cocktail party) by the same subjects. See *Task* in the main text and Supplementary Figure 1 below for details.

### Setup

Subjects were seated 50 cm in front of a 40 cm monitor (E. Systems, Inc., CA). Audio was presented diotically with a Sound Blaster Z sound card (Creative Labs, Singapore), a Scarlett 414 sound interface (Focusrite Plc, UK) and high-quality Sennheiser HD 25 headphones (Sennheiser, Germany). EEG recordings were made with a BioSemi ActiveTwo 64-channel system (BioSemi, The Netherlands) sampled at 1024 Hz; audio was re-registered with the EEG system with a ActiveTwo ERGO Opticlink (Biosemi) at the same rate. The participants adjusted sound volume to their comfortable listening level before starting the task.

### Stimuli and task

Volunteer recordings were obtained with an Olympus recorder (VN-541OC, OM Digital Solutions Corporation, Japan). Audio recordings were processed with MATLAB® (MathWorks, United States) at a sampling rate of 44100 Hz. Cosine ramps (5 ms duration) were applied at the beginning or end of the stimulus, and surrounding manual excisions made in cases of long silences and repeated filler words. All speech stimuli were marked for apparent sex and age (below or above 35 years-old). Between 2 and 4 keywords were selected per stimulus, which were limited to nouns, adverbs, or verbs in the sentence. Each sentence was also associated to at least one topic from a designated list of 13 themes.

Stimulus mixtures were created for the cocktail party task by adding two different stimuli from the database without replacement. A unique set of cocktail party stimuli was generated per participant, with equal probability of creating same- and different-sex combinations. Each cocktail party stimulus was built with a 750 ms onset asynchrony between speaker streams, and the resulting overlap time epoch marked the end of the stimulus. Stimuli were equalized in root mean squared (RMS) intensity for the overlapping epoch. All stimuli (single speaker or cocktail mixtures) were individually normalized.

In the present study, cocktail party behavioral data involves all control trials from the original studies (number of trials: 26 to 36). These trials consist of SSL phases that are unrelated to cocktail party listening phases, as they always involve different speakers, sentences, and content.

### Questionnaire and scoring

Trial questionnaires consisted of a 9-option multiple choice set with available options that were based on aspects specific to the target or background speech. Participants were asked to select three options that best matched the indicated target. Questionnaires always contained one option that was related to speaker identity, plus eight options that were related to sentence content. With equal probability, the single identity option prompted participants on vocal qualities, either of the speaker’s sex (masculine/feminine) or age (younger/older). Four additional content-related options were based on specific keywords pronounced by either or both talkers in the mix; these options were randomly sampled from the lists of keywords corresponding to the two sentences. Finally, four more content options prompted participants to report on the topic of the attended speaker. The options were randomly selected from the databases’ 12-theme list. All questionnaire options were shown in randomized order.

For scoring, any correctly chosen option always earned one point. Incorrectly chosen options deducted points as follows: a) for identity options, selecting the opposite gender or opposite age of the attended speaker resulted in a point deduction; b) for keyword options, selecting a keyword related to the masker speech resulted in a point deduction; c) for topic options, missed choices or choices related to the masker speech did not result in point deductions. Trial scores equaled the sum of point earnings and deductions across all options that were entered and validated by the participant. Validating no responses in a trial did not gain any points. A trial was deemed correct if its score was greater than zero.

### Speech spectral feature analyses

To discount the spurious spectral contributions of silent periods and weak transients in natural speech, all periods that corresponded to the signal’s envelope below a certain threshold were automatically discarded. The threshold was set per speech sample at the bottom quintile of the full broadband envelope histogram for that speech signal. Removal was performed after temporal alignment with the EEG data (see *Stimulus-response alignment and decoder estimation*, below). To obtain the broadband envelope, the raw speech signal was submitted to a bank of 28 gammatone filters with ERB-spaced centre frequencies corresponding to the 52 – 4956 Hz range. From each sub-band, the envelope was extracted by taking the magnitude value of each sample and raising it to the power of 0.6 [64]. The resulting sub-band envelopes were averaged, and this total envelope was resampled to 1024 Hz. Below-threshold epochs were discarded in corresponding periods of the envelope and in all the spectral features.

### EEG preprocessing

After preprocessing, if a head channel resulted in activity with broadband power above 3 standard deviations relative to all other head channels across the experiment session, it was removed. Single-trial presentations of solo speech stimuli were epoched, after which the variance-based automated rejection procedure was applied across single channels and epochs’ timeseries per participant. This resulted in < 1% rejected channel-epoch timeseries on average (subject range 0.09%–1.52%).

### Stimulus-response alignment and decoder estimation

Male voice presentations were selected from each participant’s EEG SSL phase data, resulting in 46trials per participant on average (range: 31 – 62 trials). For any given time sample, channel activity levels exceeding 7 standard deviations of all values in the rest of the data were set as an empty value. EEG data were temporally aligned with the acoustic feature timeseries sets (including the envelope, *f*_0_, *F*_1_-*F*_5_, and Δ*F*) by cross-correlation of the audio signal sampled by the EEG system with the corresponding raw audio original file downsampled to 1024 Hz. EEG and acoustic feature waveforms were aligned on the basis of the estimated cross-correlation delay. All data epochs when the broadband envelope signal was below a level threshold (see *Speech spectral feature analyses*, above) were removed at this point. All co-aligned EEG and stimuli epoch trials were concatenated in time. All signals were 1–8 Hz bandpass filtered with an order 2 Butterworth filter in the forward and reversed directions and resampled to 128 Hz. Prior to decoder estimation, all timeseries were transformed to z-score units.

### Statistical analysis of decoder data

As null hypothesis, slow modulations in the *i*-th auditory feature were considered to not be reliably decoded from the EEG data, which may correspond to non-positive decoder reliability (neural decoding performance) estimates across listeners. To test the alternative hypothesis, one-sided Student’s *t*-tests were run per auditory feature evaluating whether each feature modulation is reliably decodable; *p*-values were adjusted by the Holm-Bonferroni method to correct for multiple testing.

It is worthwhile noting that, in decoding studies, validation metrics may be affected by how decodability is assessed. Inspection of raw scores (empirical Π values, Supplementary Fig. 2) suggests that correlations appear generally greater than a fixed reference level of Π=0. To address generalization we have included cross-validation at the estimation stage, plus training of decoders de novo from reshuffled data. This latter step follows equal performance optimization rules as do decoders estimated from empirical data; generalization is then inferred upon greater performance for empirical-relative to reference-optimized results. Because this multi-stage approach is modeled after the statistical structure of both sampled stimuli and responses, it may impose relatively more stringent conditions than reference approaches based on models exclusively limited to the stimulus or the response.

## Supplementary Figures

**Supplementary Figure 1.**
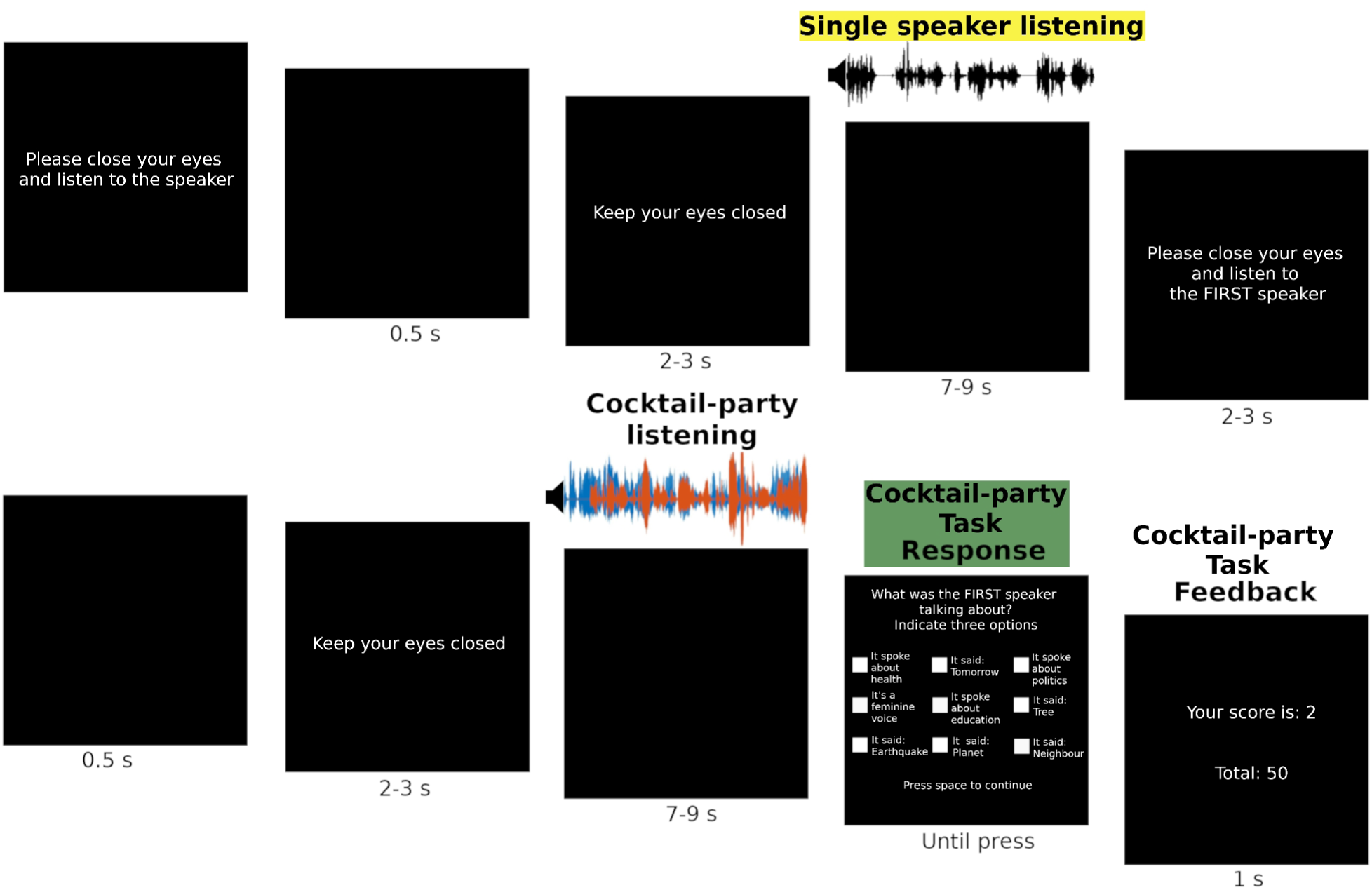
Trial design. In the first part of the trial, participants are instructed to listen to a single speaker presentation (yellow highlight) with their eyes closed. In the second part of the trial, they are given an instruction to attend to either the leading (“first”) or interjecting (“second”) talker at the upcoming cocktail party speech mixture with their eyes closed as well. After listening, a questionnaire prompts for different aspects of the indicated target speech, including speaker sex/age, topical content and specific keywords (green highlight). Trial and accumulated feedback are shown upon validation. Modified from Sánchez-Costa, et al.

**Supplementary Figure 2.**
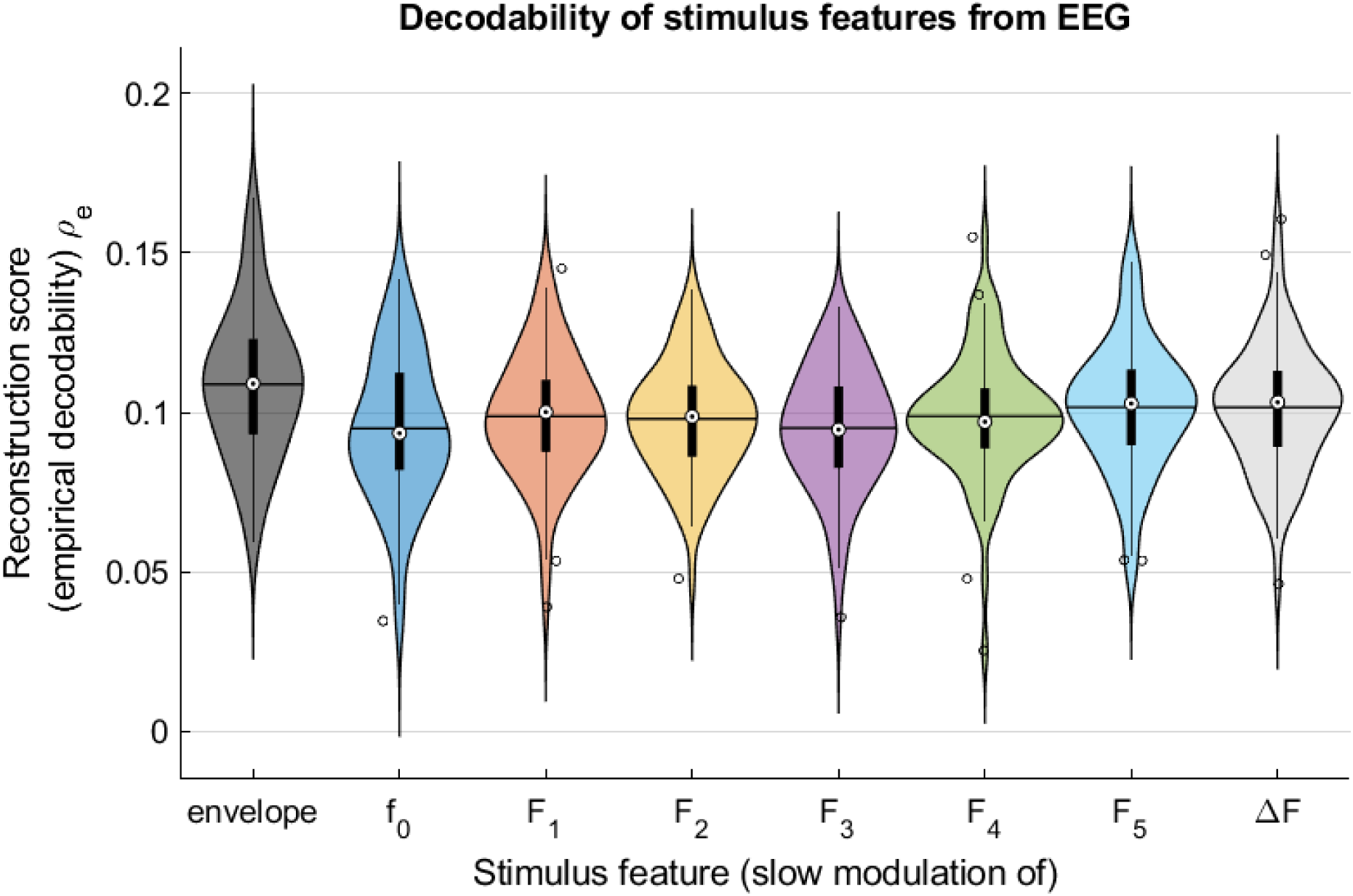
Empirical performance of EEG decoders by stimulus feature. Each plot represents the distribution across listeners (N=73) of the Spearman’s rank correlation value evaluating the ability to reconstruct empirical data, with the mean indicated by horizontal lines. Boxplots indicate distribution median, IQR as well as outlier data. Across most features, the similar reconstruction scores taken from raw decoders may illustrate the need to estimate baseline performance from reference decoders.

**Supplementary Figure 3.**
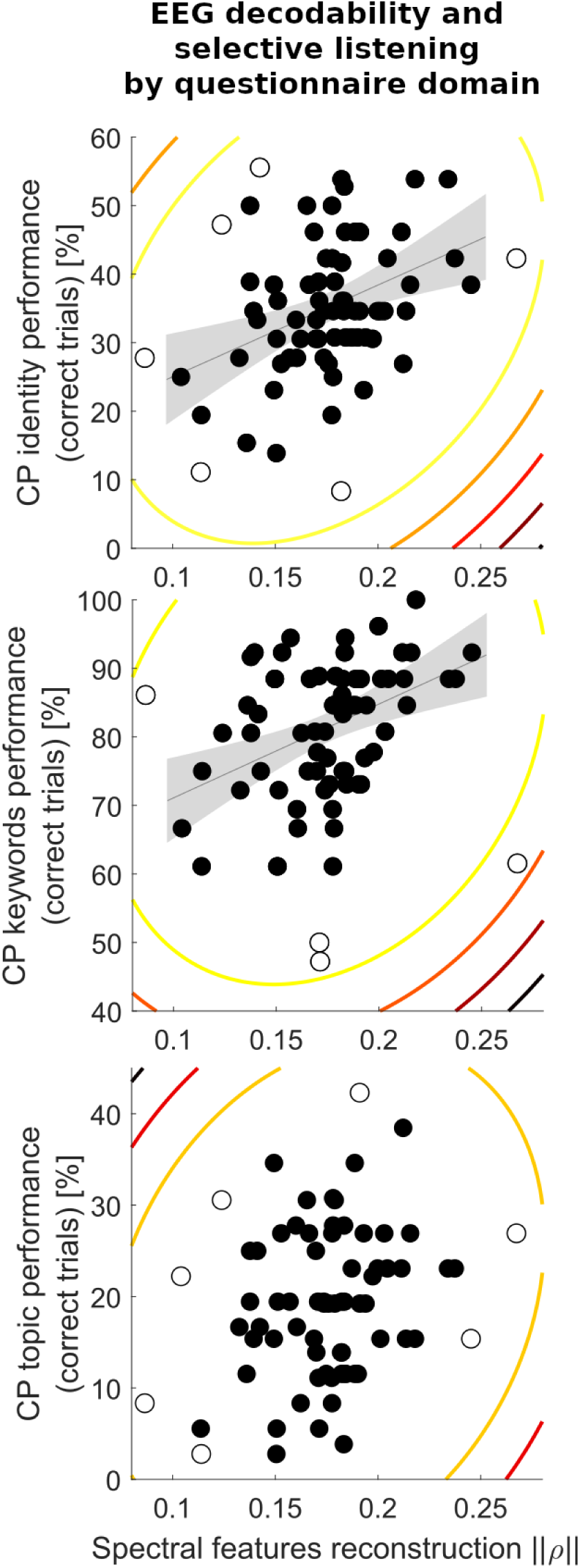
Spectral features’ reconstruction and subject performance addressed by separate aspects of the cocktail party task. (A) Listeners’ ||Π|| predicts their ability to correctly respond to the part of the questionnaire that relates to target speaker’s identity (the apparent age or sex of the voice) during selective listening. There is a significant robust correlation relationship observed between both neural and behavioral variables, Shepherd’s Pi = 0.35, *p* = 0.007. (B) Similarly, ||Π|| significantly predicts listeners’ ability to discriminate and report the words expressed by the target speaker during the ‘cocktail-party’, Shepherd’s Pi = 0.39, *p* = 0.002. (C) By contrast, listener correct responses about topic aspects of the questionnaire were not significantly predicted by || Π||, Shepherd’s Pi = 0.18, *p* = 0.319.

